# Selection and explosive growth may hamper the performance of rare variant association tests

**DOI:** 10.1101/015917

**Authors:** Lawrence H. Uricchio, John S. Witte, Ryan D. Hernandez

## Abstract

Much recent debate has focused on the role of rare variants in complex phenotypes. However, it is well known that rare alleles can only contribute a substantial proportion of the phenotypic variance when they have much larger effect sizes than common variants, which is most easily explained by natural selection constraining trait-altering alleles to low frequency. It is also plausible that demographic events will influence the genetic architecture of complex traits. Unfortunately, most rare variant association tests do not explicitly model natural selection or non-equilibrium demography. Here, we develop a novel evolutionary model of complex traits. We perform numerical calculations and simulate phenotypes under this model using inferred human demographic and selection parameters. We show that rare variants only contribute substantially to complex traits under very strong assumptions about the relationship between effect size and selection strength. We then assess the performance of state-of-the-art rare variant tests using our simulations across a broad range of model parameters. Counterintuitively, we find that statistical power is lowest when rare variants make the greatest contribution to the additive variance, and that power is substantially lower under our model than previously studied models. While many empirical studies have attempted to identify causal loci using rare variant association methods, few have reported novel associations. Some authors have interpreted this to mean that rare variants contribute little to heritability, but our results show that an alternative explanation is that rare variant tests have less power than previously estimated.

## 1 Introduction

Much recent debate among geneticists has focused on the role of rare variants in complex traits [1–4]. While there is evidence that rare variants can in some cases contribute to common diseases [5,6], it has not been established whether they explain a large proportion of the genetic variance for any traits of interest. Indeed, a number of recent papers have suggested that most of the variance is attributable to common variants of weak effect, at least for some phenotypes [7, 8]. Nonetheless, several large-scale sequencing studies focused on binary or quantitative traits are underway, and these studies will uncover many novel rare variants within their samples. Most of this variation is likely to be unrelated to the phenotype of interest, so it is imperative that population and statistical geneticists continue to make progress on methods to biologically interpret this deluge of noisy data [9].

Unfortunately, the power to detect rare causal variants with single-marker tests of association is very low, even in large samples. For this reason, several statistical methods that pool rare variants within a putatively causal locus and jointly test for their role in the phenotype have been proposed [10–18]. When pooling variants of differing effects, assessing statistical power is not a simple matter of considering all possible effect size and allele frequency combinations, because the state space of possible joint distributions is very large. Since this distribution is not known, most studies that assess the power of rare variant association tests have proposed arbitrary joint distributions that assign larger effect sizes to rare alleles than common alleles. Natural selection acting on trait altering alleles can cause an inverse relationship between allele frequency and effect size, but it is not obvious that previously studied joint distributions of effects and allele frequencies are biologically or evolutionarily plausible because most studies have not explicitly modeled natural selection (but see [19–21] for examples of studies incorporating selection).

Here, we examine the performance of popular rare variant association tests using a selection-based phenotype model. We focus primarily on one of the most popular methods, the “sequence kernel association test”, or SKAT [15]. Specifically, we assess SKAT-O [17, 18], an optimized version of SKAT that subsumes many other rare variant association tests (such as the C-alpha test [16]) as special cases. We focus our efforts on assessing this test for a broad range of parameter combinations and for a subset of our parameter space, we show that SKAT-O has better power than competing methods.

We develop a model of complex traits that explicitly captures the relationship between effect sizes and selection strength and simulate genotypes and phenotypes under the model. Following earlier studies, we show that rare variants contribute substantially to the genetic variance only under very strong model assumptions. We then apply rare variant association tests to our simulated data, and show that our selection-based model of complex phenotypes can result in dramatically lower power calculations. Moreover, power is sensitive to assumptions about recent demographic history, particularly the rate of population growth in the past several thousand years. Surprisingly, we find that the power of SKAT-O (using its default settings) is inversely proportional to the genetic variance explained by rare variants under our phenotype model, even when controlling for the genetic variance explained by the test sequence. This means that the test has the worst power when the rare variants make the greatest contribution to genetic variance. We show that power with SKAT-O can be substantially increased by adjusting its parameters, but that this approach dramatically increases false discovery rates. These results suggest that the power of rare variant association tests may have been overestimated in previous studies, and that more work is needed to develop methods for rare variant association tests (or adapt current methods) to be well powered under realistic assumptions about demographic history and recent selection.

## 2 Model

We develop a phenotype model that explicitly captures the relationship between selection strength and effect size and is a variant of models proposed in [22] and [23]. Here, we will briefly describe these two models and motivate the modifications we have made.

The model of [22] computes effect sizes *z* with *z = δs^τ^*(1 + *∊*). *δ* is −1 or 1 with equal probability, thereby allowing for both trait-increasing and trait-decreasing mutations. *τ* is an exponent that transforms the distribution of selection coefficients to allow phenotypic effects to grow faster (*τ* > 1) or slower (*τ* < 1) than selection coefficients. The central idea is that effect sizes may not have the same marginal distribution as selection coefficients, but sites with larger effects on fitness will also have larger effects on the phenotype. *∊* is a random normal variable with mean zero and variance *σ*^2^. As the variance of *∊* grows, the correlation between selection coefficient and effect size decreases.

In our study, we are concerned with how the joint distribution of effect sizes and allele frequencies impacts statistical power for discovering causal loci. It is plausible that power will depend on both the marginal distribution of effects and the relationship between effects and allele frequencies, so it is desirable to have a mechanism to hold the marginal distribution of effects constant in order to focus on the relationship between allele frequency and effects. In [23], the authors proposed a model with two selection coefficients, one strong and one weak, that has this property. With probability *ρ*, a mutation has effect size proportional to its selection coefficient, and with probability 1 – *ρ*, it has an effect size randomly sampled from the marginal distribution of selection coefficients (and then scaled by a proportionality constant). Here, we extend this model by 1) including arbitrary distributions of selection coefficients, such as a Γ-distribution of selection coefficients that was inferred for human coding regions [24] and 2) including both the *τ* and *δ* parameters from the model of [22].

Thus, our model for effect sizes *z_s_* for a site with selection coefficient *s* can be summarized as:

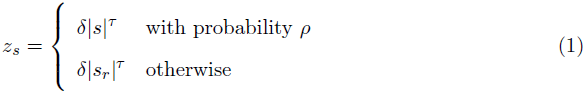

where *s_r_* is a random sample from the marginal distribution of selection coefficients.

When *ρ* = 1, we obtain exactly the model of [22]. We do not include the *∊* parameter of the original model, but this parameter was shown to have no impact on genetic architecture [22]. When *τ* = 1, we obtain the model of [23], but with arbitrary distributions of selection coefficients, and the additional possibility for causal sites to be either trait increasing or decreasing.

From an evolutionary perspective, this model captures the idea that phenotypes under direct selection will have a tight correlation between selection strength and effect size (*i.e*., high *ρ*), but the marginal distribution of effects may grow faster or slower than the distribution of selection coefficients (*i.e*., *τ* can be a value greater than or less than 1). Due to pleiotropic effects, some sites may have large selection coefficients but small effects on the phenotype (*i.e*., decreasing *ρ* allows increased emphasis on pleiotropy).

## 3 Results

### 3.1 Selection and demography impact the genetic architecture of complex traits

We investigated the impact of selection and demography on the site frequency spectrum as a function of time using numerical calculations and simulations, as well as the role of singleton variants in driving variance in genetic traits under the selection-based phenotype model of [23], as discussed in the Methods. Briefly, the selection model consists of two selection coefficients, one large (*s* = −10^−2^, 2*Ns* = −146) and one small (*s* = 2 × 10^−4^, 2*Ns* = −2.92), and two effect sizes, proportional to one or the other of the selection coefficients, as mediated by the *ρ* parameter of our model (see Methods for more detail). We focus on singleton variants because it is generally more difficult to detect extremely rare causal variants in association studies, even when multiple variants are pooled together. If there are appreciable changes in the overall contribution of singletons to the trait as a function of demographic events and model parameters, this may impact the performance of pooled tests of association.

We developed software to numerically calculate the site frequency spectrum and the genetic architecture of complex traits as a function of time using the propagation of rescaled Wright-Fisher matrices (see Methods). In the model of European demographic history proposed in [25], our numerical calculations predict that the proportion of variable sites that are singletons is strongly impacted by demographic events (solid lines, Figure 1A), and that the non-equilibrium predictions made under the model are in agreement with results from stochastic forward simulations (points, Figure 1A). In particular, expansion events increase the proportion of sites that are singletons, while contractions decrease the proportion of sites that are singletons. The vertical gray dashed line shows the current time, and the final time of the calculations is at 0.7 coalescent units (≈ 3.5 × 10^4^ years in the future). The black dashed line shows the model predictions for the neutral case (*γ* = 0, not simulated).

**Figure 1.**
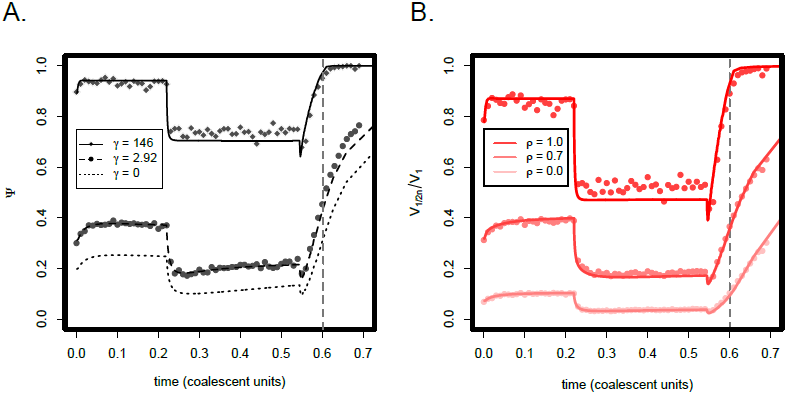
We calculated ψ, the proportion of variable sites that are singletons (A) as well as the proportion of the genetic variance in a complex trait that is due to singleton variants (B) for a sample of *n* = 100 chromosomes for the marginal European demographic history in the model of [25] and the selection-based phenotype model of [23]. The solid, dashed, and dotted lines show the results of numerical calculations under the Wright-Fisher model, whereas the points are the results of stochastic forward simulations. The vertical dashed line represents the current time, and the curves to the right of this point represent extrapolations to the future, assuming the same rate of exponential growth as was inferred in [25]. Each point represents the mean across 200 simulations.

In Figure 1B, we show the proportion of the variance in a genetic trait that is explained by singleton variants 
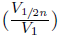
 under this model of demography, selection, and complex traits, for various values of *ρ* and *τ* = 1 (solid lines are again model predictions while the points are the results of simulations). We find that the proportion of the variance in the trait that is explained by rare variants (in this case, singletons in a sample of 100 chromosomes), is strongly impacted by demographic events, and the relationship between selection and demography. Expansions increase the proportion of variable sites that are rare, which increases the role of rare variants in the trait. Contractions have the opposite effect. When *ρ* = 1, the majority of the variance in the trait is driven by singleton variants under this model, but this proportion rapidly drops off as *ρ* decreases to 0.7 and then 0.

These results show that selection, demography, and the relationship between selection and effect sizes all impact the architecture of complex traits. Since these forces strongly impact the joint distribution of allele frequencies and effect sizes, there is reason to believe that they may also impact the performance of association tests that seek to uncover causal variation. In principle, our calculations are extensible to models of multiple populations and arbitrary distributions of selection coefficients, but at a large computational cost. We therefore build on these results by performing forward simulations under the demographic models in [25] and [26], as well as the selection model in [24].

### 3.2 Architecture of complex traits in multiple human populations under a selection-based phenotype model

We investigated the proportion of the genetic variance in a genetic trait that is due to rare variants in our model of complex phenotypes using simulations of the three human continental groups under the “growth” model in [25] and the “explosive growth” model in [26]. In Figure 2, we plot the cumulative proportion of the genetic variance due to variants under allele frequency *ω* as a function of *ω* for several different values of *ρ* with *τ* = 1 for a sample of 10^3^ individuals. Consistent with previous studies, we find that a substantial proportion of the genetic variance is attributable to rare variants only when the selection strength is very tightly correlated with the effect size. This result echoes the findings of [23], but here we have extended the previous argument by including a distribution of selection coefficients that was inferred for human coding regions [24].

**Figure 2.**
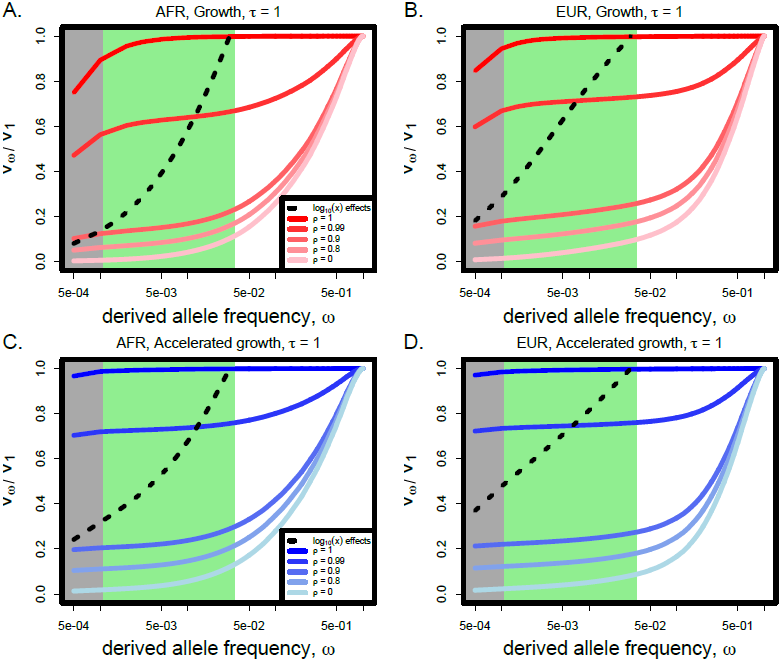
The proportion of the genetic variance 
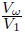
 explained by variants under allele frequency *ω* for two different demographic models of human history and two different populations with *τ* = 1.0. The gray shaded area represents alleles at very low frequency (< 10^−3^) and the green area represents moderately rare alleles (< 0.03). (*Abbreviations: EUR, European continental group; AFR, African continental group*)

We find that demography plays a subtle role in the proportion of variance explained by rare variants in the models we considered. In both Europeans and Africans, the proportion of the variance explained by rare variants increases when recent accelerated growth is included in the demographic model (*i.e*., in the model of [26]). Europeans and Africans have slightly different proportions of variance explained by rare variants for all sets of model parameters, with European demographic models tending to have slightly more variance explained by extremely rare variants than the corresponding African demographic model. We observe qualitatively similar patterns in Africans and Europeans, so we report results in Africans throughout the remaining sections, but include figures for Europeans in the Supporting Information.

Interestingly, among rare variants, the preponderance of variance explained is determined by singleton variants, and not variants at more intermediate frequencies in the sample (Figure 2). This result holds across all values of *ρ* when *τ* = 1, for both populations and both demographic models, but is more extreme in the accelerated growth model of [26]. In the black dashed lines, we plot 
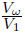
 when effect sizes are given by log_10_(*x*) for alleles with frequency *x* under 3%, which was used to generate phenotypes in [15]. Using this effect size function to generate phenotypes, much more of the variance is attributable to variants at intermediate rare frequencies.

In Figure 3, we further extend these results by considering *τ* = 0.5, meaning that effect sizes grow much more slowly than selection coefficients. Similar to the case when *τ* = 1, we find that a substantial proportion of the variance is explained by rare variants only when *ρ* ≥ 0.9. However, the demographic model subtly changes the shape of the 
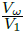
 curves, meaning that the relative proportion of the variance explained by singletons vs intermediate frequency rare variants is different under the two demographic models that we considered. Additionally, we find that nearly all of the variance due to rare variants is attributable to singleton variants when *τ* = 1, but an increasing proportion of the variance is due to moderate frequency rare variants as *τ* decreases. More variance is due to intermediate frequency rare variants in the growth model of [25] than than the accelerated growth model of [26].

**Figure 3.**
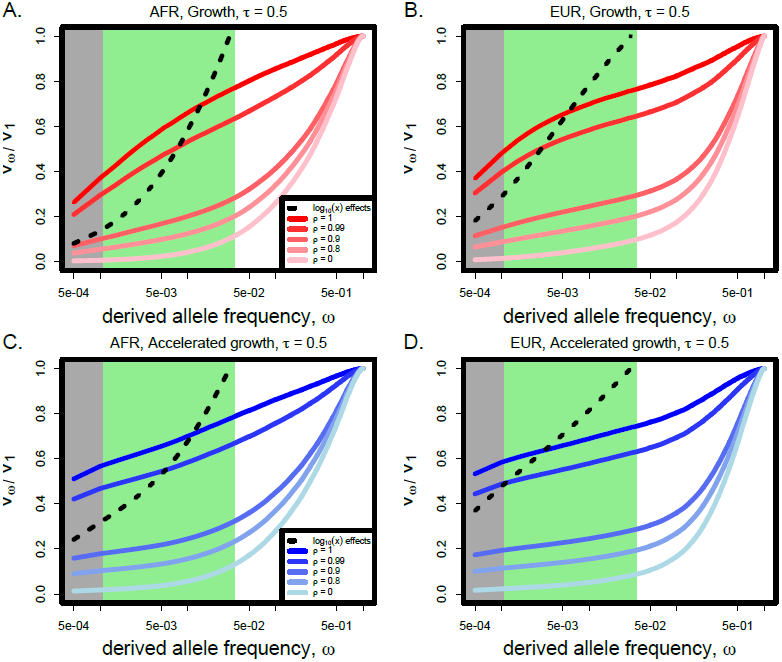
The proportion of the genetic variance 
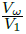
 explained by variants under allele frequency *ω* for two different demographic models of human history and two different populations with *τ* = 0.5. The gray shaded area represents alleles at very low frequency (< 10^−3^) and the green area represents moderately rare alleles (< 0.03). (*Abbreviations: EUR, European continental group; AFR, African continental group*)

These results demonstrate that the joint distribution of allele frequencies and effects are very different between our model and the log_10_(*x*) model in [15]. In particular, very-rare variants have a much larger impact relative to intermediate frequency rare variants in our model than in the previously considered model. Only when effect sizes grow much more slowly than selection coefficients (*i.e*., *τ* < 1) do intermediate rare variants play a substantial role, and recent accelerated growth decreases the relative importance of intermediate frequency rare variants. Since statistical power is a function of this joint distribution of frequencies and effects, there is reason to hypothesize that the parameters of our population-genetic model may therefore impact the power of association tests.

### 3.3 The power of conventional rare variant tests is inversely proportional to variance explained by rare variants

We investigated the power of SKAT-O – using its default settings – as a function of the proportion of variance explained by rare variants by altering the parameters *ρ* and *τ* in our simulations of complex phenotypes (Figure 4). For each of our simulations, we include 20 genes and fix the total heritability of the phenotype at *h*^2^ = 0.5, but allow the variance explained by each independent gene to vary (see Methods). To account for the variability in the overall effect on the phenotype given by each gene, we compute power as a function of variance explained.

**Figure 4.**
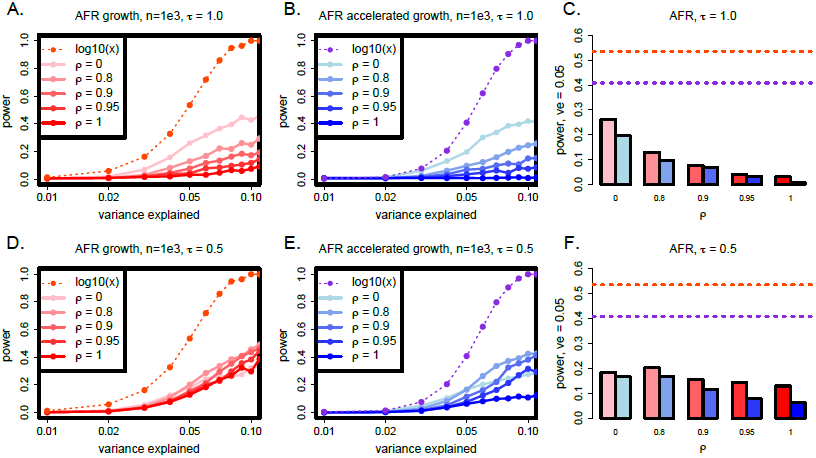
The power of SKAT-O in Africans under various effect size models, as a function of the fraction of the variance in the phenotype that is explained by the test sequence. The dashed lines show the power when the effect sizes are taken to be proportional to log_10_(*x*) for alleles at frequency *x*, while the solid lines (A,B,D,E) and bars (C,F) show results from our model (see Methods). Panels C and F aggregate data from the other panels, but specifically for variance explained equal to 0.05. The accelerated growth model of [26] is shown in blue, and growth model of [25] in red. (Abbreviations: *AFR, African continental group; ve, variance explained*).

We find that the power is substantially lower when effects are drawn from our model as opposed to effects given by log_10_(*x*) for alleles at frequency *x*, which was previously used to generate phenotypes and test the power of SKAT [15, 18]. This result holds for all model parameters and both demographic models that we considered. We also find that power is always substantially higher under the growth model of [25] than the accelerated growth model of [26]. Under the accelerated growth model, a larger proportion of the genetic variance due to rare variants is driven by very-rare variants (as opposed to intermediate frequency rare variants).

Since we compute power as a function of the fraction of the variance in the phenotype that is explained by each gene, we might expect that the power of SKAT-O would be unaffected by changing *ρ* (if SKAT-O is entirely insensitive to the joint distribution of effect sizes and allele frequencies). Alternatively, power might increase as *ρ* increases since SKAT-O is tuned to test for the contributions from rare variants, and rare variants play a greater role in the genetic variance as *ρ* increases. Counter-intuitively, we find the opposite, and the power decreases as *ρ* increases (Figure 4). This effect is most dramatic under the accelerated growth model (blue bars) and when *τ* = 1.0. When r decreases to 0.5 (Figure 4, bottom row) and intermediate rare variants play some role in the trait, the same trend exists but is less pronounced. We also observe similar overall patterns in Europeans (Figure S1).

We repeated this analysis for a subset of the parameter space for two other rare variant association tests in Europeans using the rvtests software (http://zhanxw.github.io/rvtests/), in particular CMC [11] and the test of [13]. We find that the overall trends are replicated for these tests, but that SKAT-O has better overall power (Figure S2).

In Figures S3 and S4, we investigated the impact of increasing the sample size to 2.5 × 10^3^ or 5 × 10^3^ by performing 2 × 10^3^ simulations for each parameter combination displayed. We observe only very modest increases in power for large sample sizes when *τ* = 1 and *ρ* > 0.9 (meaning rare variants explain some of the variance in the trait), but more substantial increases in power when *τ* = 0.5 and effect sizes grow more slowly than selection coefficients.

### 3.4 Shifting weight onto rare alleles boosts power but may also increase the false discovery rate

SKAT-O allows the user to supply a flexible weight-distribution over allele frequencies. The default distribution is a *β*-distribution with shape parameters 1 and 25, which gradually puts more weight on rarer alleles. As was noted in [15], the performance of SKAT may improve when a good choice of weights is made. We re-ran SKAT-O with a rare shifted weight distribution (*β*[0.5, 0.5]), which was the recommended weight distribution in [15] when only rare variants under 1% in frequency play a role in the phenotype. This resulted in substantial increases in power, although power was usually lower than under the the log_10_(*x*) effect size model with the default *β*-distribution (Figures 5 & S5). Furthermore, power is still sensitive to demography and the parameters of the phenotype model, although the trends are less dramatic than with the default weight distribution (Figures 5C,F & S5C,F).

**Figure 5.**
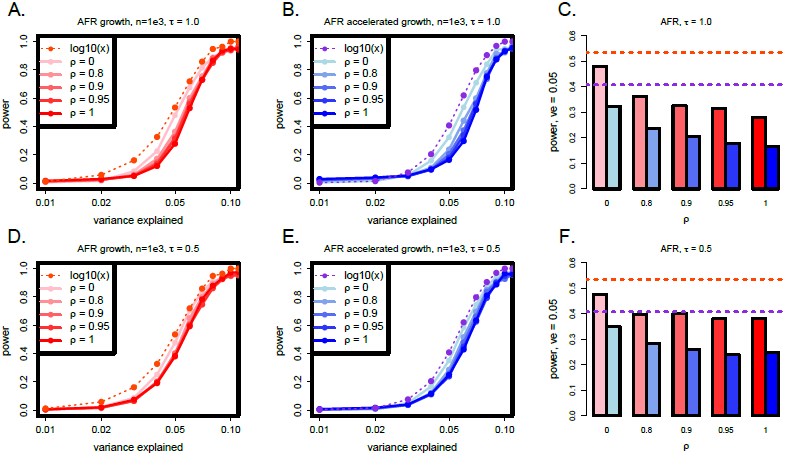
The power of SKAT-O in Africans with the weights of SKAT-O adjusted to *β*[0.5, 0.5]. The dashed lines show the power when the effect sizes are taken to be proportional to log_10_(*x*) for alleles at frequency *x* and *β*[1,25], while the solid lines (A,B,D,E) and bars (C,F) show results from our model and the adjusted weight distribution (see Methods). Panels C and F aggregate data from the other panels, but specifically for variance explained equal to 0.05. The accelerated growth model of [26] is shown in blue, and growth model of [25] in red. (Abbreviations: *AFR, African continental group; ve, variance explained*)

Unfortunately, this increase in power comes at a cost. We permuted the phenotypes for the same simulations and ran SKAT-O on the permuted data set. Under the null, we expect only 0.00025% of these tests to have p-values under 2.5 × 10^−6^, but we observe a much larger than expected fraction of p-values under this threshold. To quantify this effect, we computed precision-recall curves corresponding to a mock exome sequencing study, using the 20 genes in each simulation as the set of positive genes and 2 × 10^4^ genes with permuted phenotypes as the negative set. We find that even for very low values of recall near 0, the precision is very low, always below 10% and often below 1% when *ρ* is above 0. Furthermore, precision is much lower for the accelerated growth model than the growth model (Figures 6 & S6).

**Figure 6.**
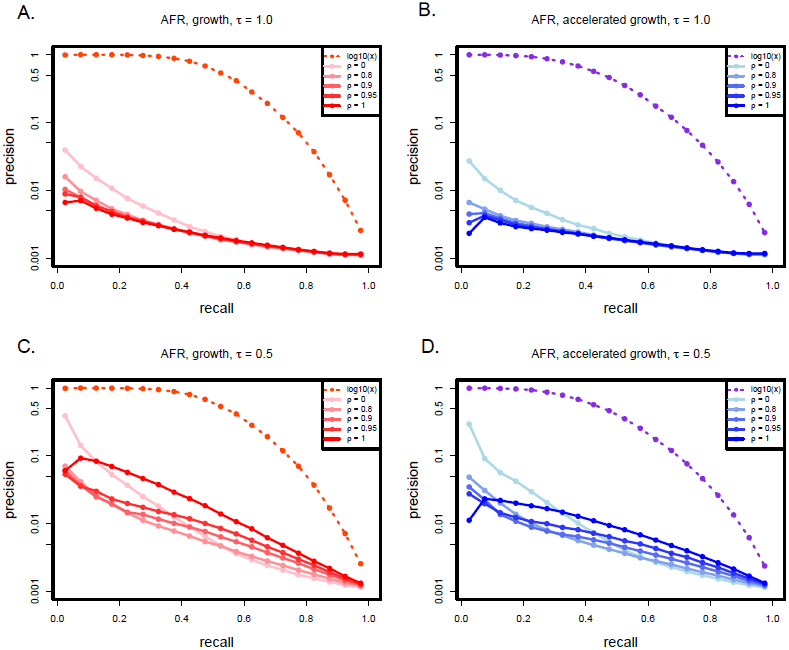
Precision-recall curves of SKAT-O for a mock exome sequencing study with rare shifted weights (*β*[0.5, 0.5]). The dashed lines, which correspond to log_10_(*x*) effects, used the default weights (*β*[1, 25]). The sample size for these simulations was 10^3^. (Abbreviations: *AFR, African continental group*)

To calculate the PR curves, we have included all 20 genes from each of our simulations as the set of positive genes. Because we have fixed *h*^2^ = 0.5, but allowed the variance explained by each gene to vary, some of the genes that we include in the positive set explain only a very small amount of the variance in the phenotype and we do not expect to be powered to discover these genes. Hence, we do not expect for any of these curves to maintain a high value of precision out to large values of recall. However, it is concerning that we observe very low precision even for values of recall close to 0, which implies that we often expect to discover many false positives even with very stringent thresholds on the test statistic.

## 4 Discussion

A great deal of research interest has focused on the problem of “missing heritability”, which refers the discrepancy between variance explained by genome-wide significant associated variants and estimates of the narrow-sense heritability of common, genetically complex phenotypes. Although there are many possible explanations for this discrepancy, one of the most popular explanations is that rare variants of large effect may make up the difference. This hypothesis has been used as motivation for a number of large-scale sequencing studies of large cohorts.

As sequencing technology has progressed to the point where very-rare (and potentially novel) variants are routinely detected in a large samples, there has been a corresponding push to develop statistical tools to detect causal rare variants. One of the most popular tools is known as SKAT-O, or the optimized sequence kernel association test [17,18]. This test is now routinely applied to large sequencing data sets.

Although it is clear that rare variant association tests are very successful at detecting associations under some phenotype models, it does not necessarily follow that previously investigated phenotype models are biologically or evolutionarily plausible. Here, we extended previous models of the relationship between selection strength and effect size to develop a model of complex phenotypes [22,23], and performed simulations under this model that include complex human demography and a previously inferred distribution of selection coefficients for human coding sequences. We showed that both genetic architecture and power calculations are quite sensitive to demography and the relationship between selection strength and effect size, and that power estimates under our model are generally substantially lower than under the previously investigated model.

A principle reason for this discrepancy is the role of very-rare variants in complex phenotypes under our model. Under our evolutionary model, when rare variants explain a substantial proportion of the genetic variance, the greatest contributions are made by extremely rare variants. When accelerated growth is included in the demographic model, very-rare variants have even larger effect sizes relative to higher frequency variants. Singleton variants are the hardest of all variants to detect since they occur only once in the data set, and non-causal singletons are ubiquitous in sequencing of large samples. We showed that some modifications to the default settings of SKAT-O can substantially improve power under our model, but these changes may also lead to an increase in false positive rates. In exome-wide studies that perform ≈ 2 × 10^4^ statistical tests, this results in very high false discovery rates, but the same does not necessarily follow for studies that perform fewer statistical tests by collapsing genes within a pathway or only testing a limited number of genes. Furthermore, we used a particular parameterization of the *β*-distribution in this study that was recommended by [15] when very-rare variants make the greatest contributions to the phenotype. It is possible that other parameterizations may strike a better balance between power and false positives.

If very-rare variants do play a substantial role in driving variance in some phenotypes at the population level, then our results may also have implications for the design of sequencing experiments, where researchers sometimes must choose between deeper sequencing and more samples. Singletons are the hardest variants to detect in sequencing data, and both power and false positive rates for calling singletons may depend strongly on the selected methods (e.g., multi-sample vs. single-sample calling). These considerations may have important implications for power of rare variant association tests if our model accurately captures the genetic architecture for some complex phenotypes of interest.

Our results demonstrate that it may be more difficult to interpret a null signal from a rare variant association test than previously appreciated, as power is highly sensitive to the parameters of evolutionary models and may be lower than was estimated in previous studies. However, though the distribution of selection coefficients that we used was inferred from human polymorphisms, it is possible that the resulting phenotype model does not accurately depict complex human phenotypes. In particular, here we focused on coding regions and quantitative phenotypes for simplicity, but some (potentially large) portion of heritability of many traits is likely to be driven by regulatory elements, which may have a very different distribution of selection coefficients [27]. Still, it is clear that *any* phenotype model that is relevant to a rare variant association test must include selection, because rare variants do not contribute substantially to phenotypic variance in the absence selection. Our evolutionary model also has implications for the distribution of phenotypes in a population. In particular, the phenotype distribution is not expected to be exactly normal because of the presence of very large effect mutations, which generate heavier tails. Models of the joint effects of selection and demography on genetic and phenotypic diversity may provide important insights into human genetic architecture and modes of selection [21–23,28–32]. In future studies, it will be advantageous to exploit these signals to compare various models of heritability (including models with and without selection) to put firmer bounds on the proportion of the genetic variance that is determined by rare variants and provide further insight into the performance of rare variant association tests.

## 5 Materials & Methods

### 5.1 Simulations of phenotypes

We simulated phenotypes under our selection-based model of complex traits, and also the allele frequency based model of [15] (the original SKAT study).

For simulations under our phenotype model, we do not have an analytical expectation for the distribution of sampled selection coefficients for the complicated three-population demographic models that we simulate here. For this reason, we use the sampled variants in any given simulation to provide a distribution on *s*. When *τ* = 1, *ρ* is also the Pearson correlation between the selection coefficient and the effect size, but this is not the case when *τ* ≠ 1. However, the interpretation that a high value of *ρ* corresponds to a high correlation between effects and a low value corresponds to a weak correlation holds across all values of *τ*. Note, we only take non-synonymous sites as causal; synonymous sites always have 0 effect in our simulations.

We also simulate phenotypes under the model of [15]. In this model, an allele with frequency *x* < 0.03 has effect size *z*(*x*) ∝ log_10_(*x*) with probability 0.05, and otherwise has an effect size of zero. Here, we take all non-synonymous sites under 3% frequency to be causal, such that the total number of causal sites are roughly comparable between simulations under our model and simulations using the effect size distribution of [15]. Note that our loci are shorter than the loci simulated by [15] because we focus on genes, but by taking all the non-synonymous low frequency variants as causal we have close to the same expected number of causal variants within the locus as in their study, though we have far fewer non-associated variants. In this sense, our simulations represent a “best-case” scenario, since a very high proportion of the sites within each causal locus are causal for the trait, though some sites have very small effect sizes.

The statistical power of association tests is a function of the fraction of the variance in the phenotype that is explained by the test sequence. For this reason, we always fix the total contribution of the test loci at a pre-specified amount (and hence any observed differences in power cannot be explained by systematic differences in the contribution of genetics to the phenotype). We simulated a polygenic model where genetic variation in the trait is driven by 20 genes, and fix the total contribution of genetics to the phenotype at 50% (*i.e*., *h*^2^ = 0.5) in Europeans. Though the choice to fix heritability at 50% is arbitrary, our results do not depend strongly on this choice since we always compute power as a function of variance explained. We have conditioned on the heritability of Europeans because the majority of genetic studies to date have been performed in populations of European descent [33]. Hence, the heritability in Africans is not guaranteed to be exactly 0.5, but can be either smaller or larger in any given simulation, depending on the effect sizes and allele frequencies of the causal variants in the sampled African sequences.

### 5.2 Calculating the impact of demographic events on genetic architecture

We investigated the impact of selection and demography on the site frequency spectrum, as well as the genetic architecture of complex traits, using numerical calculations under the Wright-Fisher model and stochastic forward simulations.

For our numerical calculations, we consider a model consisting of discrete and exponential population size changes. While our software is generalizable to other demographic models of interest, here we focused on the marginal European demographic history of the model of [25]. This model includes population size changes of magnitudes *ν* = [1.685, 0.170732, 0.47619] at times *t* = [0, 0.219178, 0.544658] (times are in coalescent units). These events correspond to an expansion in the African ancestral population, an out-of-Africa bottleneck, and a second bottleneck with the founding of Europe. Immediately after the last bottleneck event, the population grows exponentially at rate 58.4 (scaled in coalescent units). For further details on the model parameters, see [25] or [34].

To calculate the site frequency spectrum as a function of time after demographic events, we propagated Wright-Fisher transition matrices forward in time. The transition probability for a site present in *k* copies in a population of size 2*N* to *k*\* in the next generation with selection coefficient *s* is given as

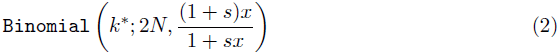

where 
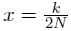
, the allele frequency of the site. Discrete changes in population size change the state space on *k*, and hence the rate at which drift happens in each subsequent generation, as well as the equilibrium proportion of variable sites present at any given frequency. However, to avoid the computational cost of changing the state space on *k*, we instead use rescale time and population size. The code we developed is implemented in Python and is freely available upon request.

We performed these calculations for two selection coefficients, *s* = −10^−2^ and *s* = −2 × 10^−4^, each with identical underlying mutation rates, exactly as in the selection-based phenotype model proposed in [23]. We assumed a human ancestral population size of 7.3×10^3^, as was inferred in [25], such that the *γ* = 2*Ns* = −146 for the large selection coefficient and *γ* = −2.92 for the small selection coefficient.

We used our code to calculate the proportion of variable sites that are present in a single copy (singletons) in a sample of 100 chromosomes for each of the selection coefficients in this model, which we denote as ψ. We also calculated the genetic variance due to singleton sites as a function of *ρ* in our phenotype model, assuming that *τ* = 1.

We performed stochastic simulations under this model of selection and demography, and sampled variants at time points from *t* = 0 to *t* = 1 (in coalescent units). We performed 100 simulations per time point. Scripts for the simulations, which were performed using `sfs_coder` [34], are available upon request of the authors.

### 5.3 Three-population forward simulations of human selection and demography

We used `sfs_coder` to perform forward simulations of human selection and demography [34]. `sfs_coder` is a python based front-end to the forward simulation software SFS_CODE [35] that includes several models of human demography and selection. The demographic models we simulated are those of [25] and [26]. Briefly, the model of [25] includes three populations, namely the African, European, and Asian continental groups. The European and Asian populations are formed by a series of bottlenecks as the human population moved out of Africa, and the model also includes recent exponential growth in the European and Asian continental groups. Migration between all pairs of populations is also included in the model. The model of [26] includes all of the above features, but also adds a second (more-recent) phase of accelerated exponential growth in the European continental group and includes recent exponential growth in the African continental group. Note that the model of [26] only inferred parameters of recent growth for Africans and Europeans, but we also simulate the Asian continental group as was inferred in [36], and as previously described in [34]. We refer to the model of [25] as the “growth” model, and the model of [26] as “accelerated growth”. We chose these two models because they both represent plausible demographic histories of human continental groups inferred from human sequence data, but propose dramatically different rates of recent expansion in Europeans and Africans, and hence generate different patterns of variation in samples. In particular, the accelerated growth of the model of [26] results in a site frequency spectrum that is skewed further towards rare variants in very large samples.

We used the selection model that was inferred by [24], which is a Γ-distribution of selection coefficients on non-synonymous sites in the human genome. Throughout each simulation, every non-synonymous mutation has a selection coefficient drawn from this distribution, while synonymous sites are neutral. In each simulation, we included 20 unlinked genes. Each gene was 1.65 × 10^3^ base pairs long, which is the mean length a gene in RefSeq. While each gene is unlinked, recombination was included within each gene. Assuming a per-base recombination rate of *ρ_r_* = 4*Nr* = 10^−3^ and that the typical gene is composed of exons and introns spanning an average of 5.115 × 10^4^ bp, we set the per-base 
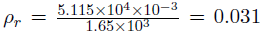
. While this does not maintain the intron/exon structure of a gene, average linkage disequilibrium across the entire gene should be maintained.

We performed between 2 × 10^3^ and 5 × 10^3^ simulations under each demographic model and sampled between 10^3^ and 5 × 10^3^ individuals (2 × 10^3^ to 10^4^ chromosomes) at the end of each simulation from the African and European populations, as noted in the text (note, as described above, each simulation includes 20 simulated genes). The number of simulations was chosen in order to obtain sufficiently small standard error around our power estimates such that parameter sets investigated are distinguishable. At the end of each simulation, we use the simulated patterns of diversity and distributions of selection coefficients to simulate phenotypes, as described above.

### 5.4 Calculating the genetic variance

We follow several earlier studies in calculating the genetic variance due to alleles at frequency *ω*, including [1] and [23]. Genetic variance *V_ω_* due to variants at or below allele frequency *ω* is given by

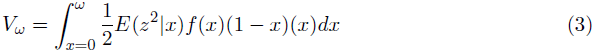

where *f*(*x*) is the site frequency spectrum, *i.e*. the proportion of sampled alleles at frequency *x*, and *E*(*z*^2^|*x*) is the mean squared effect size of variants at frequency *x*. In order to obtain an accurate measure of the site frequency spectrum and the effect sizes of variants at frequency *x*, we pool 250 simulations performed under each model, for a total of 5 × 10^3^ total simulated genes. We divide by *V*_1_, the total variance explained by genetic factors, in order to obtain a value that represents the proportion of the genetic variance explained by variants under frequency *ω*.

### 5.5 Power of SKAT-O

We obtained the SKAT-O R package from http://www.hsph.harvard.edu/skat/. We computed power as the proportion of simulations with *p*-values below 2.5 × 10^−6^. We used this threshold since our study focuses on selection in coding regions, and hence our analysis is relevant to an exome sequencing study with ≈ 2 × 10^4^ genes. 2 × 10^4^ statistical tests corresponds to a Bonferroni corrected *p*-value of 2.5 × 10^−6^. We used the default settings for SKAT-O unless otherwise stated.

### 5.6 Precision-recall curves

We calculated precision-recall curves for SKAT-O across a range of parameter values. A precision-recall curve is a plot of the precision (the fraction of positive predictions that are true positives) as a function of the recall (the fraction of true positives that have been called as positives, which is equivalent to the statistical power). A perfect precision-recall curve maintains precision of 1.0 out to a recall of 1.0. To calculate these curves, we assumed the same genetic model as above, where 20 causal genes make up the phenotype, and *h*^2^ = 0.5. In order to emulate a whole-exome sequencing experiment, we also supposed that there would be 2 × 10^4^ total sequenced genes in each experiment.

To generate the negative (*i.e*., non-associated genes), we permuted the phenotypes for all 10^5^ genes for each parameter combination and ran SKAT-O on the permuted phenotypes and genotypes. Hence, each combination of *ρ* and *τ* have their own set of 10^5^ non-associated genes.

For each of our 5 × 10^3^ sets of 20 simulated causal genes, we randomly sampled 2 × 10^4^ p-values from the set of permuted simulations to generate a distribution of null p-values. We repeated this procedure for each set of 20 genes and averaged the results to generate the precision-recall curves.

## Acknowledgments

We thank Daniel J. Balick and Raul Torres for helpful discussions, Jeff D. Wall for useful comments on an early version of the manuscript, and Dara Torgerson for supplying statistics about genes in RefSeq. This work was partially supported by a Sloan Foundation Research Fellowship (to R.D.H.), and National Institutes of Health grants 1R01HG007644 (to R.D.H), R01 CA088164 (to J.S.W.) and U01 CA127298 (to J.S.W.). L.H.U. was partially supported by an Achievement Rewards for College Scientists fellowship.

## Supporting Information

**Figure S1.**
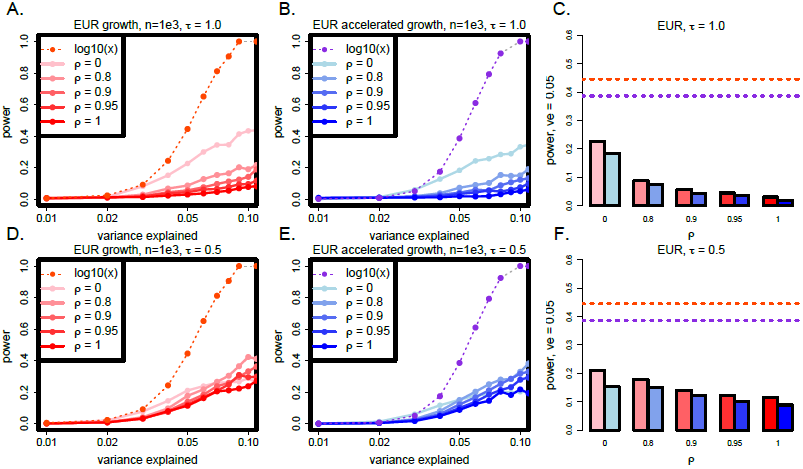
The power of SKAT-O in Europeans under various effect size models, as a function of the fraction of the variance in the phenotype that is explained by the test sequence. The dashed lines show the power when the effect sizes are taken to be proportional to log_10_(*x*) for alleles at frequency *x*, while the solid lines (A,B,D,E) and bars (C,F) show results from our model (see Methods). Panels C and F aggregate data from the other panels, but specifically for variance explained equal to 0.05. The accelerated growth model of [26] is shown in blue, and growth model of [25] in red. (Abbreviations: *EUR, European continental group; ve, variance explained*).

**Figure S2.**
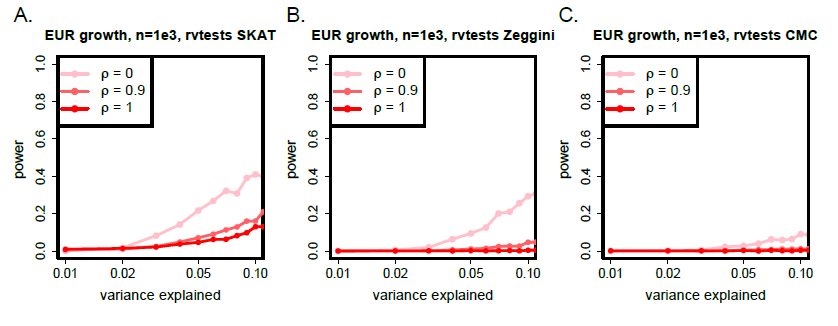
Power as a function of the variance in the phenotype that is explained by the test sequence for three different rare-variant association methods in rvtests (http://zhanxw.github.io/rvtests/). (*EUR, European continental group*)

**Figure S3.**
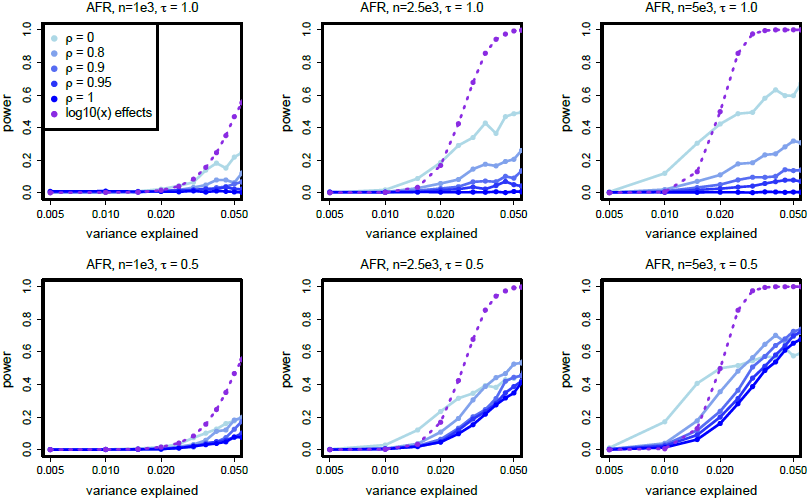
The power of SKAT-O in Africans as a function of the fraction of the variance in the phenotype explained and sample size *n*. All simulations are under the model of [26].

**Figure S4.**
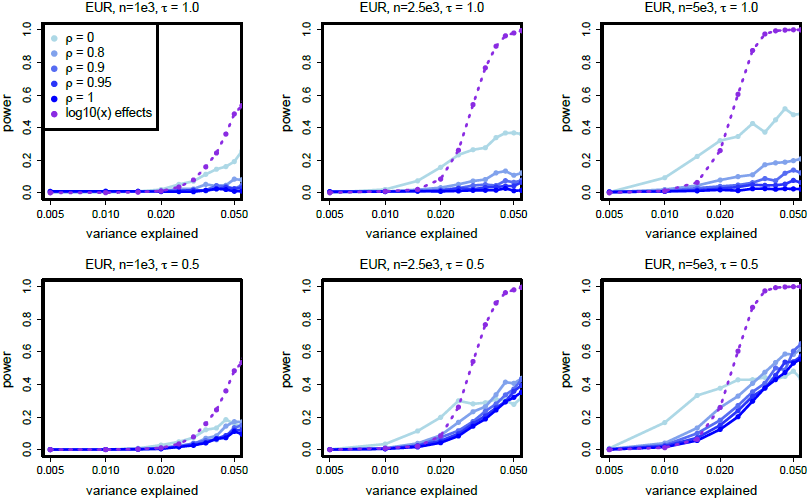
The power of SKAT-O in Europeans as a function of the fraction of the variance in the phenotype explained and sample size *n*. All simulations are under the model of [26].

**Figure S5.**
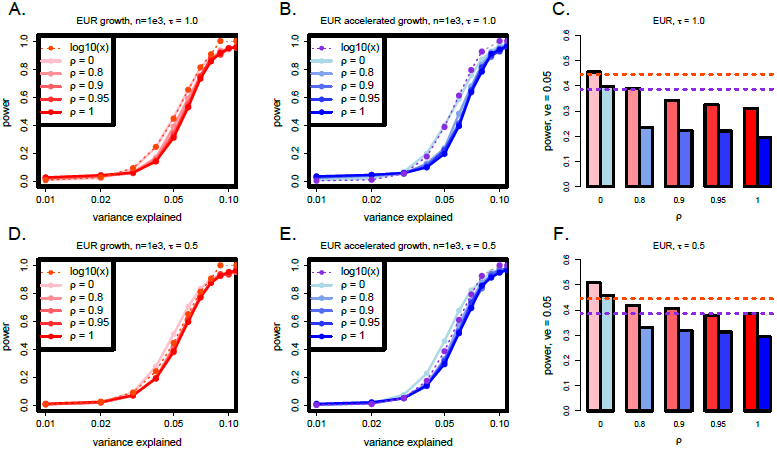
The power of SKAT-O in Europeans under various effect size models, as a function of the fraction of the variance in the phenotype explained by the test sequence, with the weights of SKAT-O adjusted to be *β*[0.5, 0.5]. The dashed lines show the power when the effect sizes are taken to be proportional to log_10_(*x*) for alleles at frequency *x* and *β*[1, 25], while the solid lines (A,B,D,E) and bars (C,F) show results from our model and the adjusted weight distribution (see Methods). Panels C and F aggregate data from the other panels, but specifically for variance explained equal to 0.05. The accelerated growth model of [26] is shown in blue, and growth model of [25] in red. (Abbreviations: *EUR, European continental group; ve, variance explained*).

**Figure S6.**
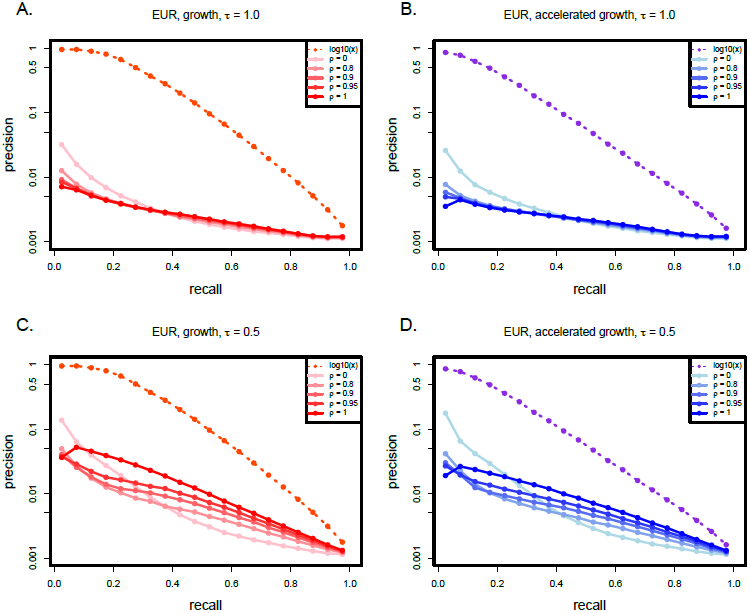
Precision-recall curves of SKAT-O for a mock exome sequencing study with rare shifted weights (*β*[0.5, 0.5]). The dashed lines, which correspond to log_10_(*x*) effects, used the default weights (*β*[1, 25]). The sample size for these simulations was 10^3^. (Abbreviations: *EUR, European continental group*)

